# Evaluation of a Deep Learning Based Approach to Computational Label Free Cell Viability Quantification

**DOI:** 10.1101/2024.08.29.610252

**Authors:** Allison Reno, Jianan Tang, Madeline Sudbeck, Precious Fe Custodio, Brandi Baldus, Elizabeth McLaughlin, Fei Peng, Hai Xiao

## Abstract

One of the most common techniques found in a cell biology or tissue engineering lab is the cytotoxicity assay. This can be performed using a variety of different dyes and stains and various protocols to result in a clear indication of dead and live cells within a culture to quantify the viability of a culture and monitor for sudden drops or increases in viability by a drug, material, viral vector, etc introduced into the culture. This assay helps cell biologists determine the health of their culture and what toxicity added substances may add to the culture and whether they are appropriate and safe to use with human cells. However, many of the dyes and stains used for this process are eventually toxic to cells, rendering the cells useless after testing and preventing real time monitoring of the same culture over a period of hours or days. Computation biology is moving cell biology towards novel and innovative techniques such as in silico labeling and dye free labeling using deep learning algorithms. In this work, we investigate whether it is feasible to train a Resnet CNN model to detect morphological changes in human cells that indicate cell death in order to classify cells as live or dead without utilizing a stain or dye. This work also aims to train one CNN model to count all cells regardless of viability status to get a total cell count, and then one CNN model that specifically identifies and counts all of the dead cells for an accurate dead and live cell total by utilizing both pieces of data to determine a general viability percentage for the culture. Additionally, this work explores the use of various image enhancements to understand if this process helps or impedes the deep learning models in their detection of total cells and dead cells.

## Introduction

Cell viability is a vital quantitative metric for understanding cytotoxicity of in vitro mammalian cell culture models for biomedical research studies. Cell viability data from these culture models can be used to further investigate the cytotoxic effects of drugs, viruses, toxins, and tumors in culture. Traditionally, methodologies such as fluorescent and colorimetric dyes and bioassays have been used as visual indicators of biological mechanisms of death, metabolism, and injury to directly quantify cell viability. This allows for direct quantification using microscopy or FACS to quantify the color change or amount of fluorescence in the sample. While the theory behind these dyes and assays is fairly simple, the application can be quite complex as it requires careful optimization and consideration of the inherent toxicity of the dyes.

The most utilized type of indicator dyes are membrane exclusion dyes. The theory behind these dyes is that an injured or dying cell will have a more permeable membrane which will allow the dye to enter inside these cells, resulting in staining them with a color or fluorescent dye so that the dead cells can be easily identified and counted^23^. This class of dyes includes commonly used dyes such as Trypan Blue, Acridine Orange/Ethidium Bromide (AO/EB live dead), Propidium Iodide (PI), and cyanine dyes (SYTOX and YO-PRO)^22^. While these dyes have been long used for viability counts and are common in many studies, the accuracy and limitations of their usage have not been thoroughly investigated until recently. For example, a recent study performed in 2020 by Chan et al. discovered that Trypan blue that enters dead and dying cells can also rupture some of those dying cells, which can artificially decrease the dead cell count resulting in a higher viability than the true value^3^. This result indicates both proof of toxicity to injured cells as well as significant accuracy issues that notably impact how these results will be interpreted going forward. The accuracy issues are from dead cells disappearing from a cell assessment because the dye ruptured the cells so that they are no longer detectable via microscopic analysis. When flow cytometry has been used to closely evaluate the accuracy of exclusion dyes Trypan Blue, Eosin Y, and AO/EB, it was found that the AO/EB assessment was much more quantitatively accurate and reliable than the other exclusion dyes and was very similar to the flow cytometry assessment^27^. This reinforces the variability in accuracy of many exclusion dyes that, while all theoretically acting on similar mechanisms, functionally get very different quantitative results. This means that all exclusion dyes cannot be relied upon as uniform tools that can be substituted for one another. Similarly, AO/EB is considered slightly less reliable than PI staining especially when used with flow cytometry because ethidium bromide is not as highly charged as propidium iodide^11^. The variability in results amongst exclusion dyes is very high which makes them a more effective qualitative method, while only the most reliable such as AO/EB and PI should be used for highly precise quantitative measurements. Even when AO/EB is utilized it must be considered potentially less effective than PI and other highly charged dyes. Additionally, most exclusion dyes mark both mammalian cells and bacterial cells, making it difficult to utilize exclusion dyes in more complex studies with coculture of several cell lines, biofilms, and/or bacterial species. This limits their use to studies with only one cell line and eliminates them from use in studies looking at the interactions between human tissues and microbial organisms such as human microbiome models.

Another common methodology for quantitative viability assessment is the use of biochemical assays that use a variety of mechanisms to differentiate between live and dead cells in culture. Examples of common bioassay mechanisms include metabolic function (MTT assays), lysosomal function (neutral red assay), ATP production, and cell adhesion^1,2^. By determining whether a vital function of a cell is intact, these assays can differentiate healthy cells from injured, dying, and dead cells within a culture. Bioassays are powerful tools that provide excellent quantitative insight into the viability of a culture. However, they have specific limitations, optimization, and confounding factors that limit their use in experiments. Often, they are used universally without taking these limitations into account. For example, with MTT assays which are a very widely used assay for cytotoxicity and cell viability, the limitations, and confounding factors of applying this assay in an experiment has been well documented in numerous studies^10^. In a study by Ghasemi et al. on the limitations and applicability of MTT to viability assessments, the study found that MTT cannot be “flippantly” applied as a straightforward single cell viability assessment as it is used in some studies. Instead, the study found that assay must be carefully optimized before being applied and that many variables must be considered in the interpretation of the results, such as cell seeding density, number of cells, amount of reagent that enters the cell, impacts of any treatments that may alter cell metabolism, impact of the phenol red in cell medium, MTT concentration, and MTT incubation time^10^. Additionally, this study highlighted that the cytotoxicity of MTT itself is still a question and an important consideration when applying MTT as an assay for studying cell viability or toxicity of a treatment, as this toxicity may slightly increase cell death above that of the baseline viability or toxicity of the treatment.

Traditional exclusion dyes and bioassays are used to quantify cell viability, but they are most likely adding toxicity above what is from treatments or experimental conditions from the dyes and reagents used for these methods. Additionally, these dyes are known to rupture dying and injured cells leading to inaccurate viability counts. Bioassays do not rupture cells, but many common bioassays, including the most used one MTT, require careful optimization of the variables that influence success of the assay and limitation of experimental factors that could interfere with the mechanisms of the assay. This limits the type of studies that these methods can be applied to as well as how universally results can be compared. Additionally, optimization of the assay creates more labor and careful experimental design for lab personnel applying the assay. All of these limitations are connected to the chemical nature of these traditional methods and the biochemical mechanisms they utilize to perform as a cell death or metabolism indicator. There is a need to create a computational method that can replicate the results of traditional biochemical methods of cell viability while eliminating the toxicity, optimization, and limited use due to the biochemical nature of the traditional viability methods. This study aims to investigate the feasibility of the application of deep learning artificial intelligence modeling for cell viability quantification as a computation method to solve the biochemical assays and dyes downfalls while replicating similar results to the traditional methods.

The rise of artificial intelligence technologies and their success in computer vision applications have inspired many researchers to classify, segment, and analyze cell images via deep neural network (DNN)-based methods^8^. Hung et al. implemented an already-proven object detection algorithm and applied it to localizing and classifying cells with different phenotypes in an image^13^. The algorithm was challenged to detect cells infected by the malaria parasite P. vivax in blood smears by detecting the subtle differences in phenotypic changes and reached 78% accuracy. However, the object detection algorithms often have to compromise between detection speed and accuracy when the objects are small and dense like cells in culture. Another popular DNN is U-net, which demonstrates the powerful capability of image segmentation. In Falk et al., a U-net was trained on pairs of co-aligned spatial light interference microscopic images and fluorescent images^8^. The trained network was able to digitally stain the live cells and dead cells, similar to a live dead biochemical staining, with over 90% accuracy. Nevertheless, U-net needs to be trained with fluorescent images as the ground truth. The toxicity introduced by fluorescent chemicals could bias the training data towards less viability and reduce the algorithms’ generality.

In this work, we studied two independent convolutional neural networks to estimate the density of all cells and dead cells respectively. The viability of the cell culture was calculated by 1-density of dead cells divided by density of all cells. This simplified approach is based on the fact that positional information of each live or dead cell is not necessary for biochemical assays and is based on the ratio of dead to live cells. These independent CNNs have been validated on test data of differing amounts of cell death induced by UV radiation. Additionally, in this work we compare a single CNN that classifies cells as live or dead with the two independent CNNs, one for dead cell quantification and one for live cell quantification, to examine whether this approach is an improvement on previous classification attempts that utilize a single CNN. Furthermore, we examine a test set of data selected from each of the experimental groups to determine if the two independent CNNs serve as a useful research tool for tracking cell viability to identify external factors impacting cell viability. The findings suggest that the use of two separate CNNs for dead and total cell quantification instead of a single classification based on positional data approach is an improvement. In addition, the two CNN approach has reasonably strong performance and, with further fine-tuning and improvements, can be a useful research tool for quantifying cell viability in culture as a label-free, computational alternative to traditional exclusion dyes and biochemical assays in order to eliminate the external toxicity, extensive labor, and tedious optimization and limitations associated with these methodologies.

## Results

The cell images data set was acquired using the procedures outlined in **Figure 1**. MDCK.1 cells were cultured in 96 well plates and then given UV treatment according to three treatment groups and a fourth mock treatment group serving as a control. The treatment groups included a high, medium, and low UV exposure amount and then an additional control well plate that sat for the same time as a UV radiation session on the countertop so that time out of the incubator was equal and would not be an external factor increasing control group viability. These groups were used in order to create a data set that theoretically would be high viability for control, somewhat high viability for the low UV exposure group, medium viability for the medium exposure group, and then very low viability for the high UV exposure group. This method created staggered values between 97-13% viability for a variable training set that would allow for evaluating the performance of the algorithm at a range of cell culture viability values. Following UV treatment, the wells were immediately imaged and then placed back into the incubator for a period of 6 hours. This incubation period of 6 hours was to allow dead and dying cells to begin to display distinct morphological characteristics of cell injury and death. Imaging was performed with brightfield phase contrast microscopy to illuminate the transparent cells without the use of dyes or staining and to allow for live cell imaging. Additionally, images later had their contrast increased further with image processing to enhance certain features for easier recognition by the models.

**Figure 1:**
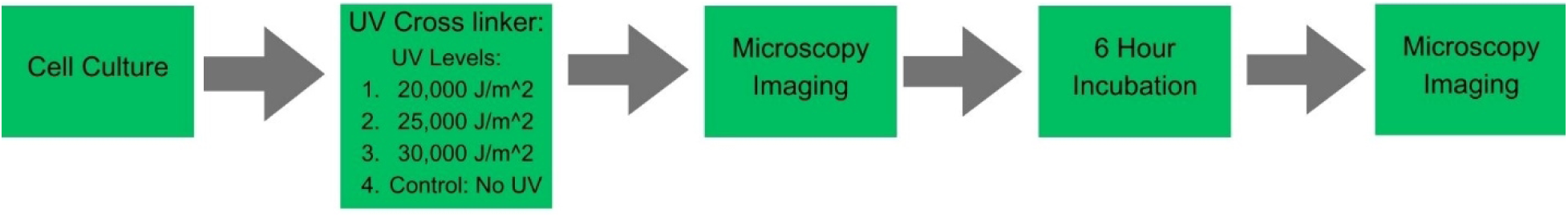

Images selected from all experimental groups were compiled and labeled as live cells only and dead cells only labeled image sets. The live and dead cell labels are combined to create the total cell data set. The selected images were cropped into 4 smaller images from each of the larger images from the original microscopy. These smaller images were labeled by lab personnel trained in cell morphology of healthy viable cells and also morphological signs of dead and injured cells. The labeling was done digitally using a VCG image annotator [https://www.robots.ox.ac.uk/~vgg/software/via/via.html]. In **Figure 2a** the lab personnel human labels can be seen in green. The green triangles in the figure mark cells labeled as live and the green Xs in the figure mark cells labeled as dead.

**Figure 2a:**
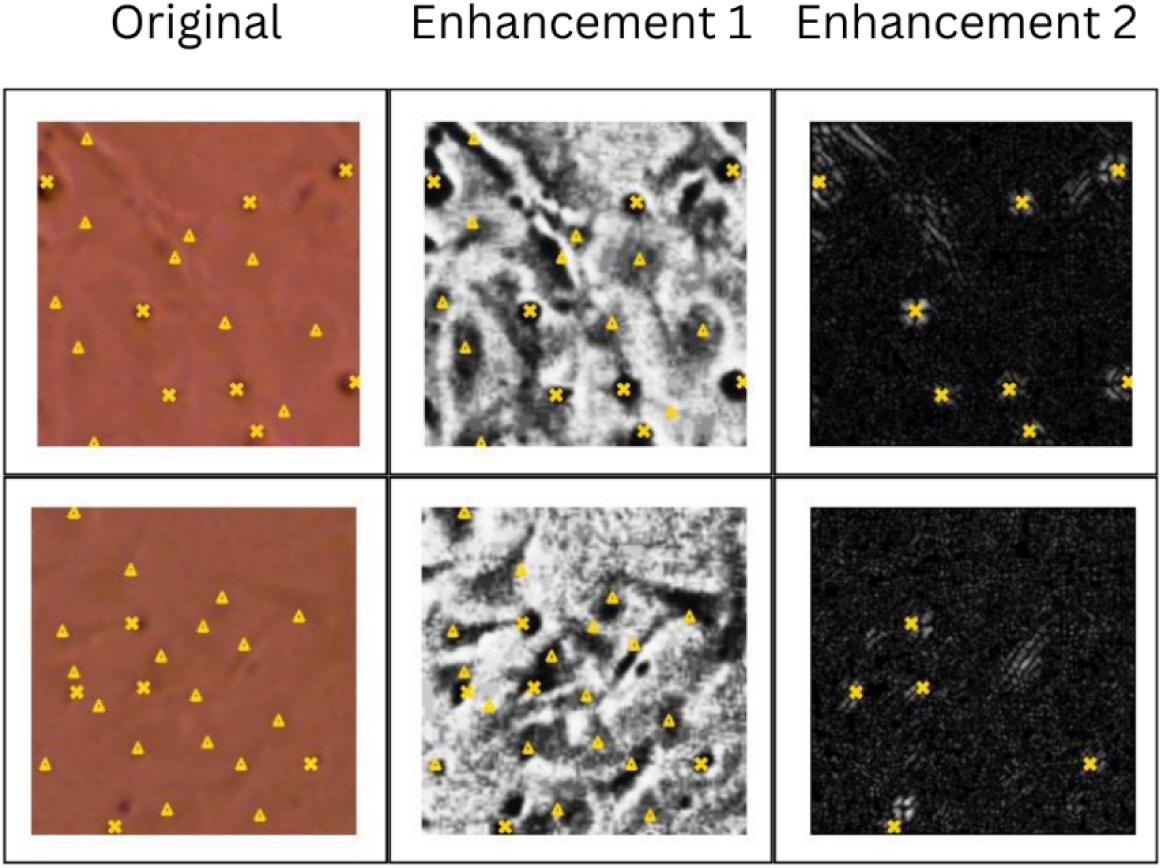

Two CNNs were trained for the automated counting; one CNN finds the dead cell count and one CNN finds the total cell count. The methodology of the two networks, one for counting total cells and one for counting dead cells, is depicted in **Figure 2b**. The original images were enhanced for more accurate detection by the CNNs by enhancing the contrast between the cells and the background. The enhancements used are shown in **Figure 2a** with a few of the raw training images and their enhancements before being used to train each model as examples.

**Figure 2b:**
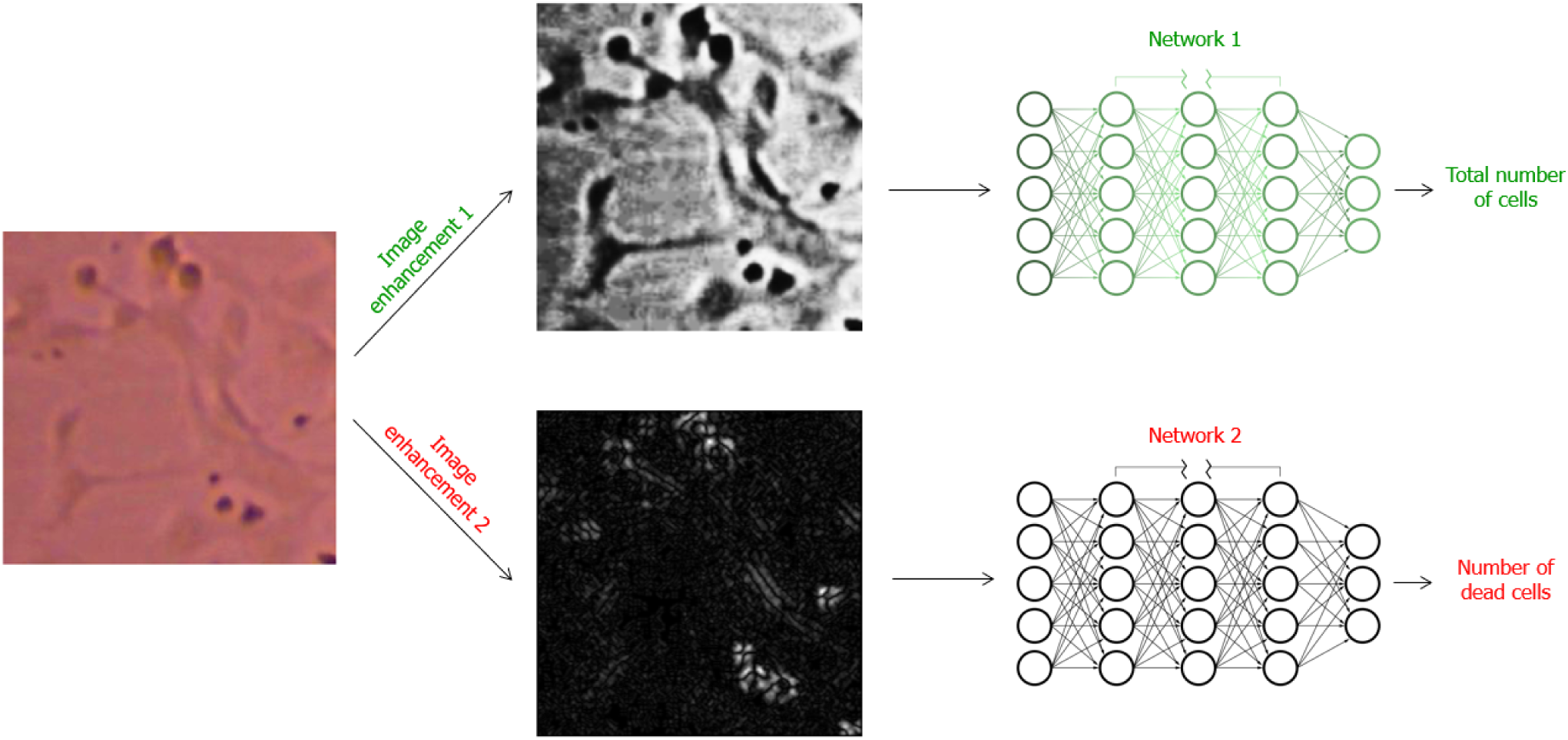

The raw image shown in Figure 2 demonstrates a color brightfield microscopy image with the lab personnel cell labeling shown in green. Enhancement 1 is the enhancement used for images before they are analyzed by the total cell CNN model. This enhancement was found to increase contrast between both live and dead cells and the background while suppressing background noise making a total cell count more easily and accurately obtained. Enhancement 2 was found to increase contrast between only the dead cells and the background while live cells do not appear as prevalent with this enhancement so both the background noise and the live cell signal is suppressed for clearer recognition of dead cells. Therefore, enhancement 2 was used on the images before they were analyzed by the dead cell CNN to increase accuracy of detection of dead cells only with this model.

The models were trained with the positional data labels in a training set of 21 number of 2048 by 1536 pixel images that were then divided into 54 smaller images tiles consisting of 224 by 224 pixels each for a total of 1134 224 by 224 image tiles. The positional labels shown in **Figure 2a** in green demonstrate the training labels that were utilized to train the CNN to recognize live and dead cells within an image and then subsequently count each detected cell for a total quantification of the total cells and dead cells within an image. Similarly, the validation set was derived from 4 of the 2048 by 1536 pixel images for a total of 216 total 224 by 224 smaller image tiles. The validation set was unlabeled, and the labels derived for this set were used for comparison purposes only as the ground truth value derivation method and not utilized for the model which was given a blank image tile. For both training and validation, the total quantification of all cells and dead cells were used as the two ground truth values to confirm the accuracy of the model’s predictions. These were derived for both the total cell and dead cell tiles by obtaining a count from the labels of each tile. Therefore, when quantifying the algorithm performance, the ground truth refers to the green labels displayed below in **Figure 2a** in the original image tiles. For clarity images were labeled by the lab personnel on the original images and then the images were enhanced after, and the labels were split into total cell and dead cell only groups and matched to the appropriate enhanced image tiles.

The individual results of the independent models showed that the total cell counter performance tended to have a slight trend of predicting above the ground truth value for the total cell count. In contrast the dead cell CNN model performance showed a trend towards under counting the dead cell value. The graphs of the results for the total cell and dead cell counter CNNs can be found in **Figures 4a and 4b**, respectively. The viability percentage results from combining the results of the two CNNs to predict viability and is depicted in a regression graph in comparison to the ground truth derived from the labeled data in **Figure 4c**. The results from the dead cell counter CNN model and the total cell counter CNN model were combined into one viability score that is a percentage out of 100 by dividing the dead cell count value by the total cell count value, multiplying by 100 to convert to a percentage, and then subtracting this from 100% to find the total viable cell percentage. The percentage for the human labeled ground truth was derived from live cell count divided by total cell count and then multiplied by 100 to convert to a percentage. The results in **Figure 4c** demonstrate that when the total cell counter CNN and dead cell counter CNN are combined the overall result tends to moderately overpredict the viability in comparison to the ground truth viability score. Because the dead cell counter underpredicts and the total cell counter slightly overpredicts in comparison to the ground truth for both values this would result in an overestimation of the number of live cells and as a result the viability percentage is also overestimated by this combined effect. This is further demonstrated by directly comparing the ground truth viability percentage to the one derived from the cell counter CNNs, as shown in **Figure 4c**, where the viability percentage is overestimated which confirms that the total cell counter’s slight overprediction combined with the dead cell counter’s slight underprediction results in a moderate overprediction of viability due to the cumulative effect of combining these errors.

**Figure 4:**
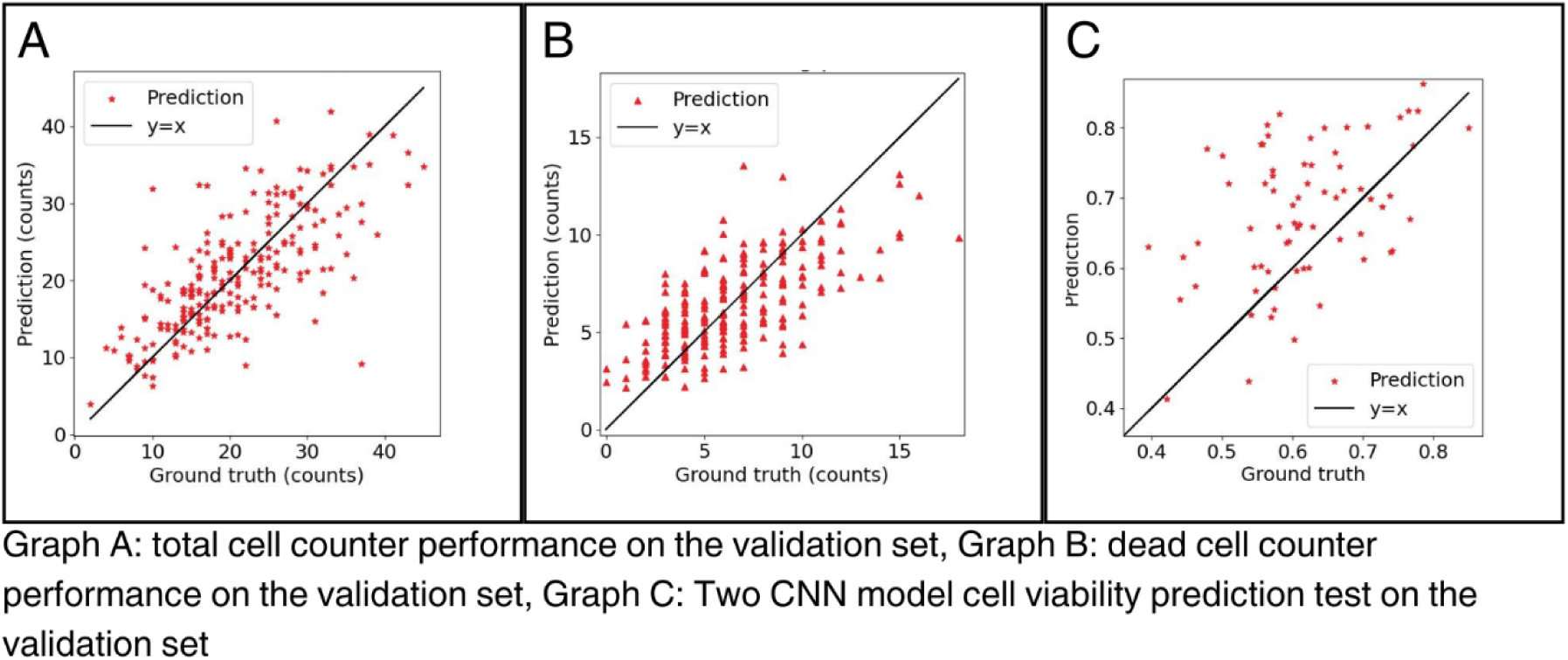

## Discussion

During the course of this study, several different approaches to determining cell culture viability using deep learning CNN models were attempted and compared in order to determine the best approach. The goal was to determine the simplest approach that also had an acceptable level of accuracy and to add further, more complex steps or processes as needed if results were not an acceptable level of accuracy until a model was found that was the minimal amount of complexity while meeting the desired accuracy. As a result, the first approach taken was the simplest and that was to use a single CNN with no image enhancements and to train this model with labels of the image’s overall viability. The goal was to have a model that would be shown a single unenhanced image and give back a total viability percentage prediction for that image. This direct approach assumed that the model over time would learn to detect differences in the total image and in the individual cells in the image that correspond to the different lab personnel viability scores being used as ground truth in order to directly result in an accurate prediction of cell culture viability percentage. The results of this single CNN approach can be found in **Figure 5a**. The results of this approach showed that the single CNN model very consistently tended to overpredict the viability when compared to the ground truth lab personnel labeling. It was hypothesized that maybe the algorithm was detecting real viable cells that the lab personnel overlooked since labeling is a very subjective technique and some variance is to be expected. However, upon investigating individual examples of the images it became apparent, especially with very low viability examples, that more real live cells did not exist or did not exist to the extent of the overestimation and that the algorithm had a tendency to predict higher than accurate values especially concerning middle to low viability ground truth images.

**Figure 5:**
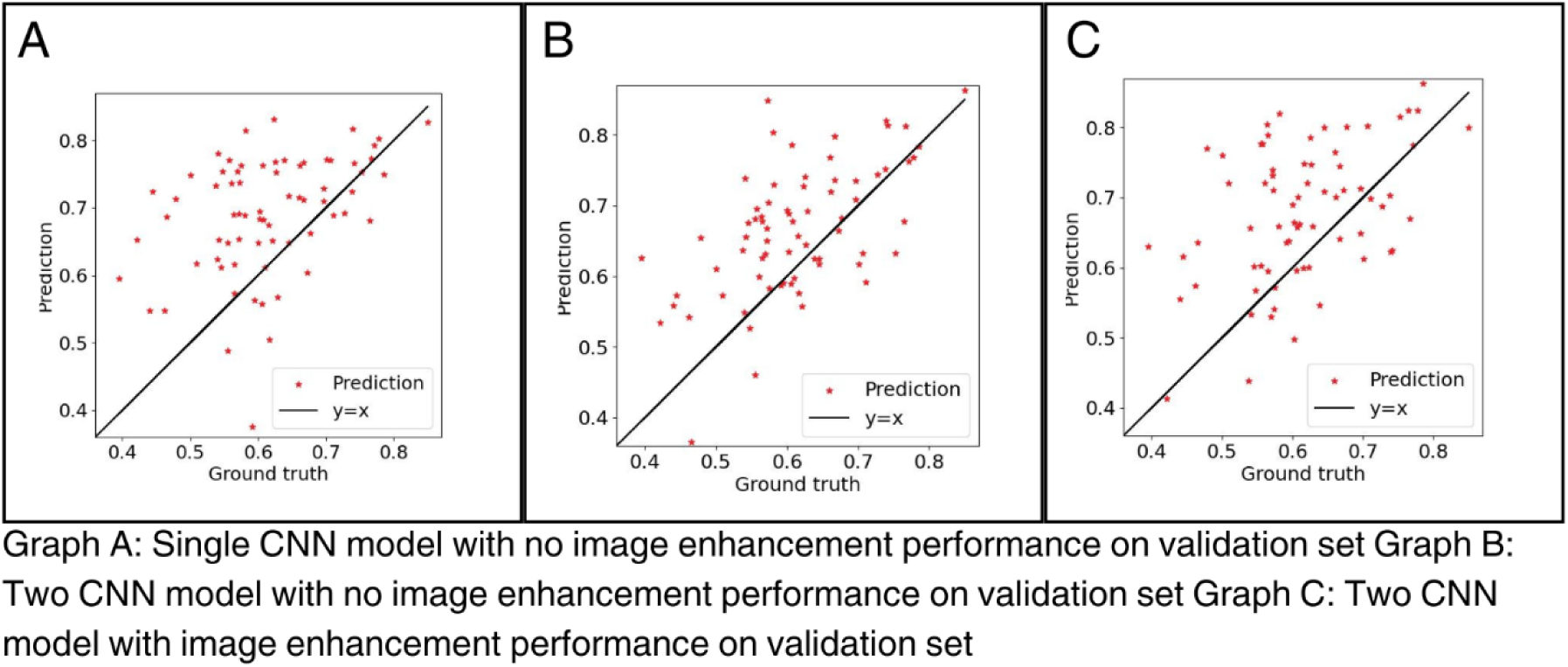

When reflecting on the shortcomings of the single CNN approach, we hypothesized that a more accurate and thorough labeling and training process was needed than viability estimations based on observation of the overall image and morphological observations of the cell health within that image. The most precise routine lab-based methodology for deriving viability is to do a direct count using exclusion dyes or morphological analysis and then using the live and dead cell counts to then calculate a viability percentage for the image. The only method more accurate than this would be to perform live/dead fluorescent staining in order to derive the live and dead cell count which is far less routine and more complex than the use of exclusion dyes or morphological analysis. Considering that the approach was maximum simplicity, the technique of morphological analysis of black and white cell images was chosen. In order to create training data sets of morphological analysis by lab personnel, the lab personnel labeled the live and dead cells in separate data sets using VCG annotator software. Then one data set was used to train one CNN, the live cell counter, and the other set of data was used to train the other CNN, the dead cell counter. This basic approach to using a 2 CNN model and the new approach to the labels used for the ground truth is shown in **Figure 5b**. Surprisingly, these changes did not increase the accuracy of the deep learning model very much only slightly correcting the overprediction and randomness of the previous approach. It was quickly apparent that this approach was a bit too simple and was allowing the CNNs to pick up on extraneous factors in the image such as brightness of the image, amount and shape of artifacts in the background of the image, clumping and clustering of the live and/or dead cells, etc that lead to results that were very far from ground truth values. While this approach did not bring a significant improvement in accuracy it did demonstrate the model was starting to narrow in on specific features of the images with this approach, even if those features were incorrect. As a result, we hypothesized that if we utilized image enhancement to eliminate extraneous factors in the images shown to the model, we might derive a model more accurate than the original approach by improving and fine-tuning the performance of this two CNN approach.

The third model attempted for this study is the final model that the results section highlights and displays the full breakdown of results from and the enhancements used for this model can be found in **Figure 2a**. This model, shown in **Figure 5c**, uses a very similar approach to the second model, but with this model while the lab personnel still label unaltered images, the images that are given to the model are altered to enhance the presence of the type of cell the counter is supposed to detect. Additionally, instead of a live cell counter, it was found that making a total cell counter was more effective as the model had particular issues with distinguishing a live set with consistently accurate results. For the enhancements the contrast was increased significantly for Enhancement 1, shown in **Figure 2a**, which eliminates background noise while increasing the contrast between the cells and the background so that it is easy to detect and visualize the edges of both dead and live cells. For Enhancement 2, Sobel kernel is applied on the image tiles. Because dead cells have larger contrast edges around them, Sobel filter can effectively enhance the presence of dead cells. In the processed image, the live cells are no longer prevalent while dead cells are presented as white dots. Using this approach, the total cell and dead cell counter were trained with their corresponding enhanced images and the combined total cell count and the dead cell count respectively in order to obtain the resulting predictions. This approach was the most successful outcome of any of the CNN models and was determined to not exceed the level of complexity desired for the study, so it was selected as the ideal appropriate model and further trained and validated with a validation set in order to fully analyze the results of this final deep learning model.

While the two CNN with enhancement model became the deep learning model selected as most successful for this study, it still has some shortcomings and requires evaluation of where it excels and still lacks in performance. The overall trend of this model is still to overpredict the validation. The breakdown of the two separate CNNs feeding into this model can be found in the results section in **Figures 4a and 4b**. Where the dead counter CNN underpredicts and the total cell counter overpredicts which combined results in a more increased overprediction of viability. For the model to be improved both the CNNs feeding into the viability result would need to be improved. The dead cell counter would need to detect a larger percentage of the true dead cells and the total cell counter would need to be refined more to stop detecting background noise as real cells. However, in comparison to the model using only one CNN seen in **Figure 5a** the overprediction is much less and is significantly improved by using the two CNNs with the enhanced images trained to estimate based on cell morphology and not overall viability appearance of the whole image. This model is still the best choice out of the 3 techniques tested and a much simpler approach than other attempts to utilize AI quantification of unstained cells, but it is important to acknowledge the areas where the model still needs refinement and further fine-tuning for future increased performance. The performance would most likely need to be increased before this could be used as a reliable and accurate research tool.

When reviewing the performance of the model, there are still times when the selected model for this study, the two CNNs with enhanced images, fails to recognize subtle morphological indications of both dead and live cells. Specific image examples, found in **Figure 6**, have been pulled from the validation set in order to analyze the performance and give examples of errors upon close analysis of the model. One large flaw of the model is that the dead cell counter CNN is not sensitive to the large range of sizes and morphologies that dead cells can exhibit during the process of cell injury and death. Example 5 serves as a great demonstration of why this phenomenon is occurring and what biological marker is being underdetected by this model. In example 5, the human count is much higher than the algorithm count which fails to recognize many of the very small dead cells. Cell death can take many different forms and there are many mechanisms outside of necrosis, apoptosis, and autophagy^9^. However, in terms of morphological analysis mostly what is seen is either cell death via swelling, classified broadly as necrosis, and cell death via shrinkage, classified broadly as apoptosis^25^. While the specific mechanism is important biologically for certain analyses it is not relevant for broadly quantifying and observing morphological markers of cell death^7^, so generally the human technicians were trained to look for signs like circular appearance, haloing effect indicating pulling up from the plate, signs of necrosis like swelling, and signs of apoptosis such as shrinking. As seen in example 5, there are many very small dead cells that are caught and counted by the human analysis. It appears though that a likely explanation for the large underrepresentation of dead cells by the algorithm count in example 5 can be explained by the algorithm mostly looking for signs of necrosis and ignoring cells with apoptosis, that are very small and hard to detect even by human technicians, as background artifacts and not a real biological phenomena of cell death shrinkage. The image enhancement technique applied may also be filtering out very small dead cells and contributing to this phenomenon which is further proven by the comparison of the total and dead enhancements in example 5 where the total cell enhancement makes these tiny cells much more clear and easy to visualize leading to an accurate count the same as the human analysis while the dead cell enhancement captures the larger and medium sized dead cells, but does not as clearly represent these very small dead cells. That is one trade-off of the use of enhancements as part of the model methodology. While using enhancements will help the algorithm more accurately tune out background noise, it may also serve to filter out or further amplify background noise by selecting what is the signal and amplifying that in a very coarse way without the ability for fine-tuning and selection. Overall, this large underestimation in dead cells does lead to a difference in the overall viability by about 14% which is a very significant amount of error. Similarly, in example 2, a similar phenomenon occurs where the total count is very accurate, but in this example the very small dead cells are ignored by the model and also filtered out by the enhancement leading to a significant underrepresentation of the dead cells. In this case, the total cell count is also underestimated leading to a much smaller difference in viability with about a 6-7% overestimation of the total viability.

**Figure 6:**
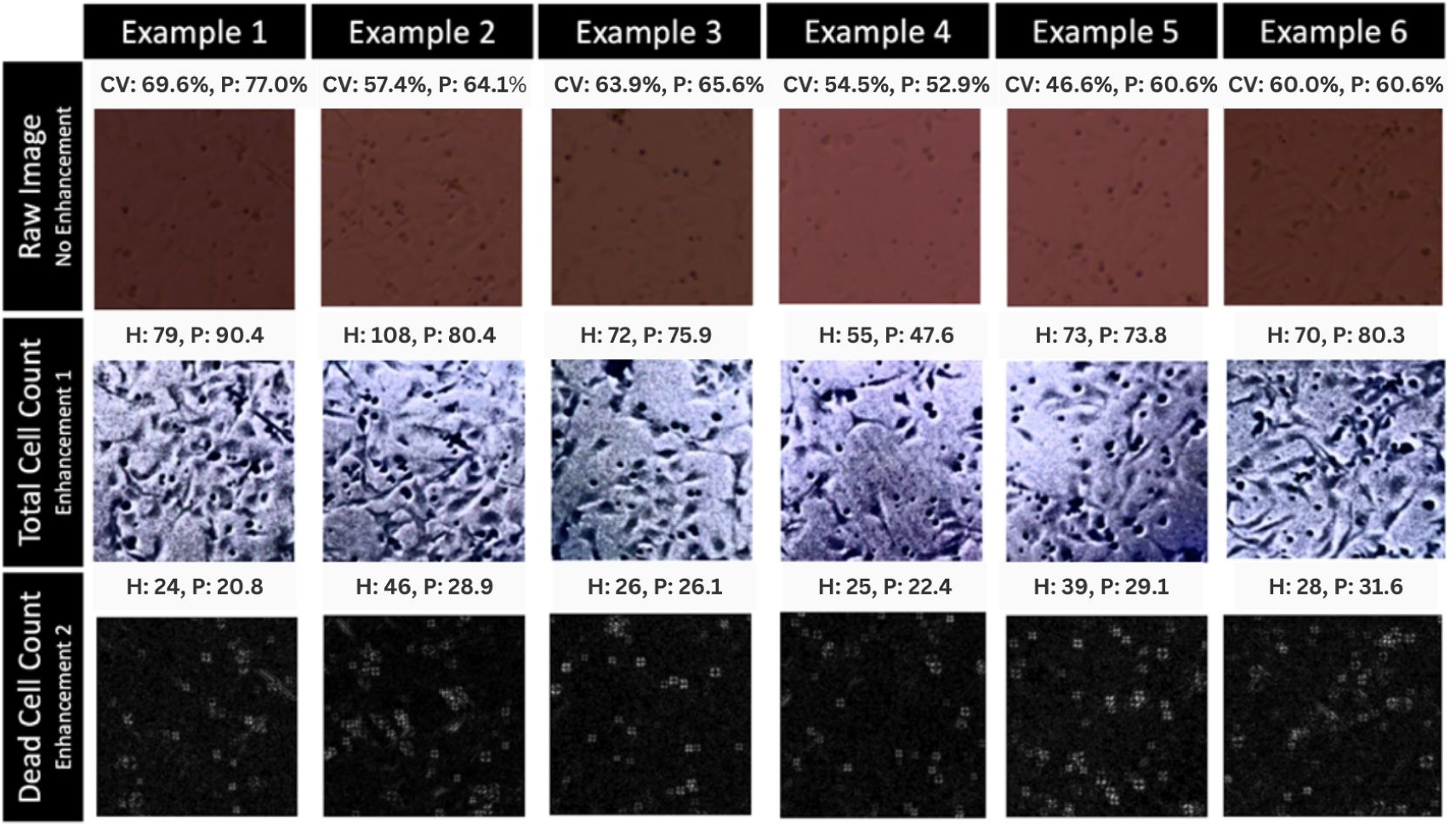

Another trend observed in the model was that in some cases the total cell count was being overpredicted. This is seen in examples 1, 3, and 6 with example 1 and 6 as especially good examples of how this impacted the overall viability prediction. In all 3 of these cases it is likely that the filter applied to increase contrast and therefore, make very subtle cell edges more pronounced making cell detection and quantification much easier, was likely also making background artifacts more prevalent. By increasing contrast with a blanket filter on the total image there is not much selection that can be done and as a result some background artifacts are amplified and made to look even more like cells which likely is the cause behind overestimated total cell counts. For example 6, this effect is canceled out by the previously discussed trend of underestimation of dead cells and ends up with the correct viability score. However, for many images including example 1 and 3 the dead cell count was fairly accurate in these cases and then as a result the number of live cells is overestimated. The total viability percentage, in the case of example 1, was overestimated by about 7% which is a very significant figure. In example 3 the difference is more subtle, and the overestimation is only about 1.7%. Additionally, the dead cell count is accurate for this example, so the overall percentage is not exaggerated by a dead cell count error and is only an error attributed to the total cell counter CNN.

These examples of fairly significant differences between the predicted viability percentage and the human label derived viability percentage highlights the need to further investigate the use of enhancements and filters to increase accuracy of quantification models. Most importantly, it is necessary to examine if they can be fine-tuned to better select for real artifacts or whether methods not using enhancement might better represent real values and not lead to models being trained to recognize false data and rule it in as a real representation of a cell. While the model performance overall does appear to be successful, taking a closer look at these examples certainly highlights that significant improvements would be needed to apply this model as a real-world research tool for accurate automated label-free cell quantification.

More rarely, in some examples the total cell count would underpredict the total count leading to substantial decrease in viability from the true value. Example 4 highlights this phenomenon where the total cell count and dead cell count are both underpredicted with the total count being much more substantially underpredicted. This led to a rare error of an underestimation of cell viability by 1.6%. This type of error, however, was found to be extremely rare within the validation set and was most likely due to human error of the human personnel labeling artifacts in the image as real cells that were then excluded by the filter applied for the total cell count. Since this was not a significant, prominent, or very high-impact error that made very little impact on the overall performance of the model, it is safe to conclude this is just a result of human error that is unavoidable when using unlabeled data. No changes would need to be made or recommended based on this result as when it did happen the underestimation of the viability was typically quite small and this was a rare occurrence for the model to underestimate instead of overestimate or accurately predict viability. This phenomenon is well under the acceptable threshold to not be concerned and not make any changes to specifically address this type of error although it is important to acknowledge a few of these cases do exist within the validation set.

Overall, this study presents a look at three different approaches to utilizing deep learning CNN modeling to try to rapidly quantify cell viability using dye free label free cell images. Models were compared, and their approaches evaluated in order to examine different approaches that can be used to improve accuracy while maintaining a simple, not time-consuming, low-cost approach and also preserving the goal of rapid, accurate quantification as a useful future research tool for cell culture laboratories. The highest performance model was selected and then analyzed with a validation set to more closely study this most successful model as an example of both the advantages and disadvantages of the methodology used to derive the model. Ultimately the use of image enhancement when viewed more broadly seems like an obvious choice as it greatly improved the accuracy of the model performance results. However, upon closer inspection of specific examples from the validation set it becomes clear that utilizing this technique also has its drawbacks and does contribute to some general trends of error within the data that explain the overall result of slight overprediction of the cell viability percentage. A more selective method of image enhancement or alteration that allows for greater fine tuning while still helping to decrease signals from the background noise is necessary to achieve the very high levels of accuracy such a model would require to be utilized in real world research scenarios. In the future we hope to identify methods to help increase the performance of the model to be able to start applying the use of the model to some small real world research scenarios to better understand the performance when applied to a real cytotoxicity cell culture study.

## Methods

### Method section 1: Cell Culture and Cell Experimentation

#### Cell Culture

The cell line used for the experiment was MDCK.1 (ATCC, USA) The cells were cultured in Minimum Essential Medium (MEM; 11095-080, Gibco, USA) supplemented with 10% fetal bovine serum (FBS; 26140-079, Gibco, USA) at 37°C with 5% CO2.

The cells were cultured with Minimum Essential Medium (MEM; 11095-080, Gibco, USA) supplemented with 10% fetal bovine serum (FBS; 26140-079, Gibco, USA) once the cells reached 70% confluency. For the subculture protocols, the adherent cells were removed from their original flasks using 0.25% Trypsin-EDTA (1X) (25200-072, Gibco, USA). Once trypsinized, the suspension was neutralized using growth medium. The cells were then placed in a conical tube and centrifuged for 7 minutes at 1500 rpm. The preexisting medium and trypsin were removed and replaced with fresh growth medium. A cell count was carried out using a Millipore Scepter 3.0 with the 40 μm filter to determine seeding density. Additional fresh medium is added to the new vessels periodically to support cell growth, and the cells were incubated at 37°C with 5% CO2. The subculturing was done a minimal number of times and then the cells were frozen as MDCK.1 cell stocks until each stock was needed for experimental cultures so that all cells used were low passage cells that maintained the qualities of the original cell line.

For the experiment, cell stocks from the MDCK.1 cell line were pulled from the -80°C freezer and seeded into four separate 96-well plates. The 96 well plates were seeded at a seeding density of approximately 100,000 cells per well. Culture plates 1 and 2 were from the same frozen cell stock and culture plates 3 and 4 were from the same frozen cell stock and both cell stocks used had identical cell culture conditions, passage number, and cold storage conditions and length of time. After the cells were split into their respective 96-well plates, the plates were incubated at 37°C with 5% CO2 for 72 Hours until they reached 80% confluency.

#### Data Sets

For the experiment plates 1-4 were assigned to experimental groups. Plate 1 was assigned as the control plate. Plate 2 was assigned to radiation group 1 also referred to as group 1 during data analysis. Plate 3 was assigned to radiation group 2 also referred to as group 2. Plate 4 was assigned to radiation group 3 also referred to as group 3.

#### UV Radiation Exposure

Once the cells reached the desired confluency, the plates were separated into a control and three experimental groups. For detailed protocols on mock treatment of the control group and the timing of radiating cell plates without overexposure to a cold environment see the experimental protocol section. Medium was removed from the treatment and mock treatment plates to prevent red phenol dye interference with radiation treatment. A Fisherbrand UV crosslinker with five 254 nm UV C lamps was used to deliver even and controlled UV exposure to induce controlled cell death at varying levels. The control group received mock treatment of medium removal and then being placed on the counter for the same amount of time as group 1 radiation exposure. Group 1 was radiated with 20,000 J/m2 of UV C light exposure for two minutes. Group 2 was radiated with 25,000 J/m2 of UV C light exposure. Group 3 was radiated with 30,000 J of UV C light exposure. Following irradiation, the phosphate buffered saline was removed and replaced with growth medium.

#### Imaging

Phase contrast imaging was performed with EVOS XL Core Imaging System with a 10x objective with a numerical aperture of 0.25 with built in LED illumination of the imaging system. Images were captured using the built in 3.1 megapixels color camera in the XL Core imaging system. The images captured were 2048x1536 pixels JPG image files.

Cell plates were imaged on a schedule to get precise time data. All groups and control were imaged at hour 0 and 6 following irradiation treatment and were imaged with phosphate buffered saline in each well to allow clear visualization on the color camera. Once timed live cell imaging concluded, a Trypan Blue membrane exclusion dye was introduced to act as a cell viability stain and images were taken immediately after introduction of the dye to prevent excessive effects of toxicity from exposure to the dye in the imaging results. Cell viability staining was done with Trypan Blue membrane exclusion dye diluted with phosphate buffered saline to a 1:20 ratio of dye to PBS. Then after the entire plate was stained, the stain was immediately removed and replaced with PBS for clear visualization of the dye on the color camera.

#### Experimental Protocol

Four 96 well plates of MDCK.1 cells were prepared from frozen stock and cultured for 72 hours to reach 80% confluency on the day of the experiment.

1. Cell plates 1 and 2 were removed from the incubator and the 300 µL/well of growth medium was removed and replaced with 100 µL/well of phosphate buffered saline to prepare the plates for radiation.
2. Plate 2 was radiated using a UV crosslinker. Meanwhile, Plate 1 was placed on a clean counter in the same laboratory during the radiation of plate 2 in order to have the same environmental exposure to serve as a mock treatment group. A timer for 6 hours was started at the completion of the treatment and mock treatment.
3. Cell plates 1 and 2 were imaged consecutively after plate 1 completed mock treatment and plate 2 completed radiation exposure.
4. 100 µL/well PBS was removed from cell plates 1 and 2 and replaced with 300 µL/well of growth medium.
5. Cell plates 1 and 2 were placed back into the incubator to await the second round of imaging after 6 hrs.
6. Cell plates 3 and 4 were removed from the incubator and the 300 μL/well of growth medium was removed and replaced with 100 µL/well of PBS to prepare for irradiation.
7. Cell Plates 3 and 4 were radiated consecutively using a UV crosslinker with plate 3 being treated first and plate 4 being treated second during plate 3 imaging and 6-hour timers were set after both treatments.
8. Cell plates 3 and 4 were imaged consecutively to obtain 0 hr images after both plates finished radiation.
9. 100 µL/well of PBS was removed from plates 3 and 4 and replaced with 300 µL/well of growth medium.
10. Cell plates 3 and 4 were placed back into the incubator to await the second round of imaging after 6 hrs.
11. Six hours after plates 1 and 2 were placed back into the incubator, they were removed individually, and the growth medium was removed and replaced with 100μL/well PBS to obtain hour 6 images.
12. Following 6-hour imaging for plates 1 and 2, plate 1 was stained by removing 100 μL/well of PBS and adding 100 µL/well of a 1/20 dilution of trypan blue with PBS and then subsequently removing and replacing with 100 μL/well PBS.
13. Plate 1 was imaged to obtain stained image data set.
14. Plate 2 was stained as specified in step 12 and imaged to obtain stained image data set.
15. Six hours after plates 3 and 4 were placed back into the incubator, they were removed individually to obtain hour 6 images.
16. After completion of timed imaging for both plates, plate 3 was stained by removing 100 μL/well PBS and adding 100 µL/well of a 1/20 dilution of trypan blue with PBS and then subsequently removing and replacing with 100 μL/well PBS.
17. Plate 3 was imaged to obtain stained image data set.
18. Plate 4 was stained as specified in step 16 and imaged to obtain stained image data set.

### Method section 2: Deep Learning Model Development

#### Generating Labeled Training Set

To create a training data set of images for the deep learning model, images were labeled by lab personnel. The images obtained by the EVOS XL Core Imaging System were annotated using a VCG image annotator software to create two labeled training sets: one training set of live cell labels and another set of dead cell labels. Lab personnel analyzed images for morphological signs of cell viability to label the “viable” and “nonviable” cells to create the two data sets. Signs of cell viability included signs that the cell was attached to the plate such as elongation, lighter coloring, no presence of a halo, and no spherical appearance. Signs of nonviable cells were signs that a cell was detaching or detached from the tissue culture vessel including spherical appearance, halo effect around the cell, dark color, small size, and lack of elongation. Cell images were uploaded to the VCG image annotator and for the live set the viable cells were identified and labeled by marking with a small circle marker in the middle of the cell. Once all viable cells were marked the JSON file containing the coordinates of the labels was saved. The unviable set was labeled in the same way and once all nonviable cells were marked with a circle marker in the middle of the cell then the JSON file containing the coordinates of the annotations was saved.

#### Model Development

Both the algorithm for predicting the total number of total cells and the number of dead cells share the same network structure. The network structure is based on EfficientNet^26^, which is a commonly used backbone structure for computer vision tasks. The top layers of EfficientNet is replaced by a global average polling layer followed by a fully connected layer. The fully connected layer only has one neuron to output a single scalar representing the number of cells in the input image. Because the training set is relatively small, transfer learning technique^28^ ^i^s applied to strengthen the feature extraction capability and generalizability of the algorithms. The networks inherit all the weights from a EfficientNet trained on ImageNet^6^, which is a large-scale image dataset that contains millions of images of different objects, such as cats, dogs, and cars. Although these images are largely different from the cell culture images, the pretrained EfficientNet is able to learn to extract common features such as corners and edges, which is widely believed to be beneficial for various kinds of biomedical tasks^5,15,17^. The networks are then finetuned on the train sets described above, without freezing any weights, to learn the distinctive features of cells.

## Acknowledgements

Krista Henrie and Rebecca Levis for help with editing and generation of figures and data tables.

## Contributions

A.R., J.T., and H.X. conceived the idea. A.R. designed the cell experiment, collected data, generated figures, and conducted data analysis. A.R. and M.S. prepared cell cultures and performed cell imaging. J. T. designed the computation parts of the experiment, performed the image filtering for enhanced cell counting, and performed the deep learning testing. A.R., M.S., P.F.C., B.B., and E.M. performed labeling of cell images for deep learning training sets. A.R., J.T., M.S., and P.F.C. wrote the manuscript with input from all authors. H.X., F.P, and J.T. supervised the project.

